# Down-regulation of LRIG1 by miR-20a modulates gastric cancer multidrug resistance

**DOI:** 10.1101/186403

**Authors:** Lin Zhou, Xiaowei Li, Fan Zhou, Zhi’an Jin, Di Chen, Pin Wang, Shu Zhang, Yuzheng Zhuge, Yulong Shang, Xiaoping Zou

**Author notes:** These authors contribute equally to this paper (Lin Zhou, Xiaowei Li, Fan Zhou, Zhi’an Jin). To whom correspondence should be addressed: Dr. Yulong Shang, Pro. Xiaoping Zou, Mailing address: Department of Gastroenterology, Nanjing Drum Tower Hospital, The Affiliated Hospital of Nanjing University Medical School, No.321 Zhongshan Road, Nanjing 210008, China.

## Abstract

Multidrug resistance (MDR) significantly restricts the clinical efficacy of gastric cancer (GC) chemotherapy, and it is critical to search novel targets to predict and overcome MDR. Leucine-rich repeats and immunoglobulin-like domains 1 (LRIG1) has been proved to be correlated with drug resistance in several cancers. The present study revealed that LRIG1 was overexpressed in chemo-sensitive GC tissues and decreased expression of LRIG1 predicted poor survival in GC patients. We observed that up-regulation of LRIG1 enhanced chemo-sensitivity in GC cells. Interestingly, miR-20a, which was overexpressed in GC MDR cell lines and tissues, was identified to regulate LRIG1 expression by directly targeting its 3′untranslated region. We also found that inhibition of miR-20a suppressed GC MDR, and up-regulation showed opposite effects. Moreover, we demonstrated that the miR-20a/LRIG1 axis regulated GC cell MDR through EGFR mediated PI3K/AKT and MAPK/ERK signaling pathways. Finally, LRIG1 expression in human GC tissues is inversely correlated with miR-20a and EGFR. Taken together, the newly identified miR-20a/LRIG1/EGFR link provides insight into the MDR process of GC, and targeting this axis represents a novel potential therapeutic strategy to block GC chemo-resistance.

## Introduction

Gastric cancer (GC) remains the second leading cause of cancer-related deaths worldwide in recent years, with a low 5-year survival rate especially for advanced patients(Siegel et al., 2017; Torre et al., 2015). Chemotherapy is the main approach used to manage GC patients of advanced stages(Van Cutsem et al., 2016). However, chemotherapeutic approaches often fail in clinical practice, due to intrinsic or acquired drug resistance, particularly multidrug resistance (MDR)(Van Cutsem et al., 2016). Although the mechanisms underlying MDR of GC have been extensively investigated, the key determinants are still largely unknown.

Leucine-rich repeats and immunoglobulin-like domains 1 (LRIG1), a transmembrane protein, is a pan-ERBB negative regulator that inhibits EGFR (epidermal growth factor receptor) signaling by accelerating receptor internalization and degradation in a c-CBLe dependent manner(Gur et al., 2004; Powell et al., 2012; Wang et al., 2013). Substantial evidence indicates that LRIG1 is involved in tumorigenesis and functions as a tumor suppressor(Wang et al., 2013). Low expression of LRIG1 has been associated with poor prognosis in breast(Krig et al., 2011; Thompson et al., 2014), oropharyngeal(Lindquist et al., 2014) and nasopharyngeal(Sheu et al., 2014) cancers. It was reported that overexpression of LRIG1 suppresses malignant glioma cell growth by attenuating EGFR activity and negatively regulates the oncogenic EGF receptor mutant EGFRvIII(Stutz et al., 2008). Moreover, LRIG1 down-regulation attenuates TNFα (tumor necrosis factor α) gene expression and confers resistance to Smac mimetics in breast and ovarian cancers(Bai et al., 2012). Recently, it was demonstrated that LRIG1 could reverse MDR in glioblastoma, by negatively regulating EGFR and suppressing the expression of ABCB1 (ATP-binding cassette, sub-family B member 1) and ABCG2 (ATP-binding cassette, sub-family G, member 2)(Liu et al., 2015). However, the function of LRIG1 in GC MDR remains unclear, and needs to be elucidated.

MicroRNAs (miRNAs) are small, evolutionarily conserved, non-coding RNAs of 18–25 nucleotides in length, which bind to the 3′-untranslated regions (3’-UTR) of target genes, leading to either target mRNA degradation or protein translation reduction(Bartel, 2009; Yates et al., 2013). It was estimated that approximately 30% of human genes and most genetic pathways are regulated through miRNAs(Bartel, 2009; Yates et al., 2013). As a negative regulator at the posttranscriptional level, miRNA is an important mechanism regulating gene expression(Bartel, 2009; Yates et al., 2013). What’ more, mounting evidence has shown miRNAs are involved in GC development and progression, including MDR(Ishimoto et al., 2016; Riquelme et al., 2016; Song and Ajani, 2013). We thus speculate that dysregulation of LRIG1’s regulatory miRNAs might be a main source that causes its down-regulation in MDR of GC.

In the present study, we showed that LRIG1 was overexpressed in chemo-sensitive GC tissues compared with chemo-resistant ones, and decreased expression of LRIG1 predicted poor survival. We further observed that up-regulation of LRIG1 enhanced chemo-sensitivity to various drugs in GC cells. In addition, we demonstrated that miR-20a, which was overexpressed in GC MDR cell lines and tissues, regulated LRIG1 expression by directly targeting its 3′-UTR. We also found that inhibition of miR-20a suppressed GC MDR, and up-regulation of miR-20a showed opposite effects. Moreover, the miR-20a/LRIG1 axis regulated GC cell MDR through EGFR mediated PI3K/AKT (PI3K: phosphatidylinositol 3 kinase; AKT: protein kinase B) and MAPK/ERK (MAPK: mitogen-activated protein kinase; ERK: extracellular signal-regulated kinase) signaling.

## Results

### Decreased LRIG1 expression is associated with poor prognosis and chemo-resistance in GC

To clarify the expression and clinical significance of LRIG1 in GC, we analysed the data from *Oncomine* ( https://www.oncomine.org/resource/login.html) and *Kaplan Meier plotter* ( http://www.kmplot.com/analysis/index.php?p=background). It was found that LRIG1 was significantly down-regulated in GC compared to normal gastric tissues in 4 independent cohorts (Figure 1A). Furthermore, individuals with lower LRIG1 expression exhibited reduced overall survival in a cohort containing 876 GC cases, and decreased progression free survival in a cohort of 641 GC patients (Figure 1B). These findings indicated that LRIG1 might serve as a biomarker in GC and lower expression of LRIG1 is associated with poor prognosis.

**Figure 1.**
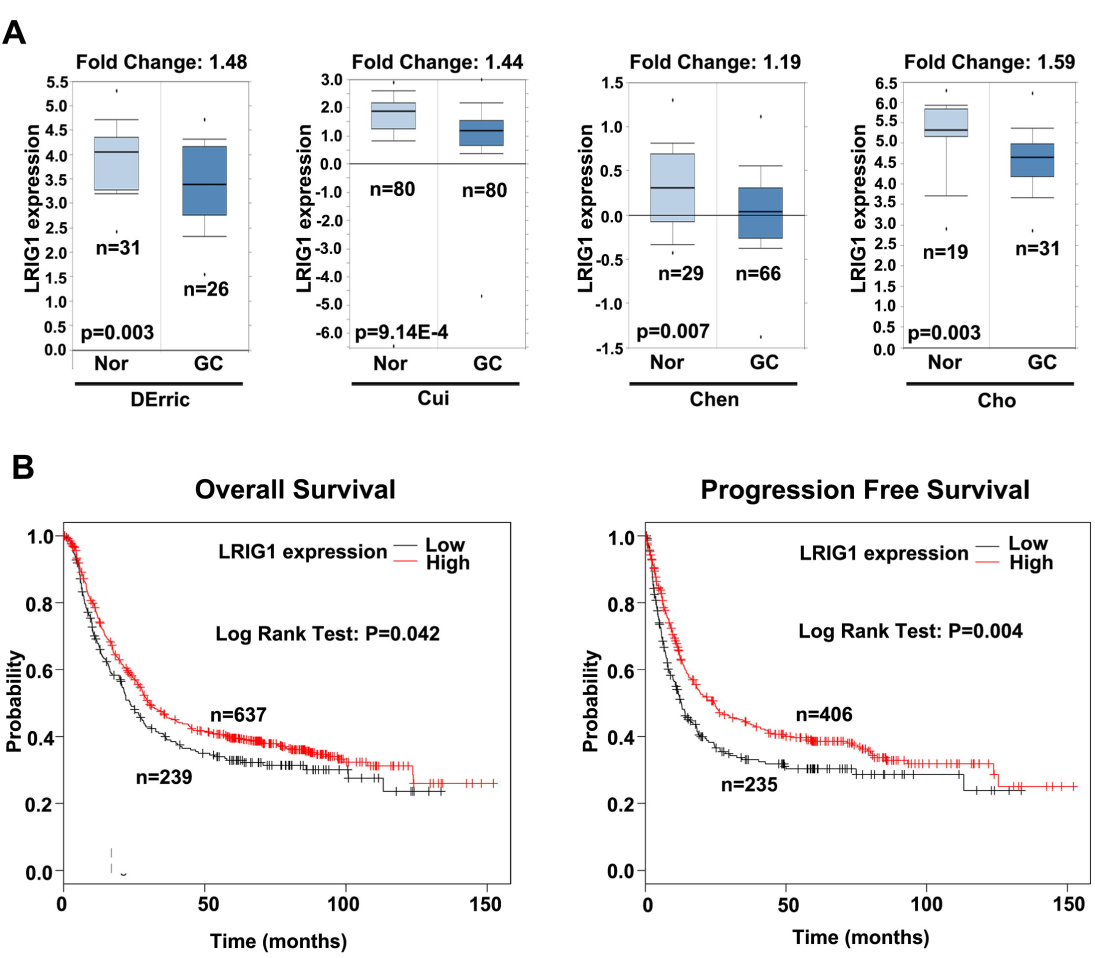
LRIG1 is down-regulated in GC tissues and associated with poor prognosis. (A) The comparison of LRIG1 expression between cancerous and normal gastric tissues from *Oncomine*. Nor, normal gastric tissues; GC, gastric cancer. (B) The association of LRIG1 with overall survival and progression free survival in GC from *Kaplan Meier plotter*.

To explore the role of LRIG1 in GC drug resistance, the expression of LRIG1 in GC tissues displaying different chemo-sensitivities was examined. The response to chemotherapy of 47 GC individuals was estimated according to the mRECIST (modified Response Evaluation Criteria in Solid Tumors) criteria, and 13 patients who were evaluated as “CR (complete response)” or “PR (partial response)” were regarded as sensitive to chemotherapy. Immunohistochemistry (IHC) and qRT-PCR analyses showed that LRIG1 expression was more frequently present in the chemo-sensitive GC cases than in the chemo-resistant ones (Figure 2A, 2B and Table 1). Moreover, IHC results further confirmed that LRIG1 expression was down-regulated in GC tissues, compared with adjacent nontumor (Supplementary Table 1). To further characterize the function of LRIG1 in GC chemo-resistance, we measured the expression of LRIG1 in GC chemo-resistant cell lines — SGC7901/ADR and SGC7901/VCR, and their parental cell line SGC7901. Western blotting and qRT-PCR analyses demonstrated that LRIG1 expression was markedly lower in SGC7901/ADR and SGC7901/VCR, compared with SGC7901 (Figure 2C and 2D). Thus, these findings confirmed that the low expression of LRIG1 correlates with enhanced GC drug resistance.

**Figure 2.**
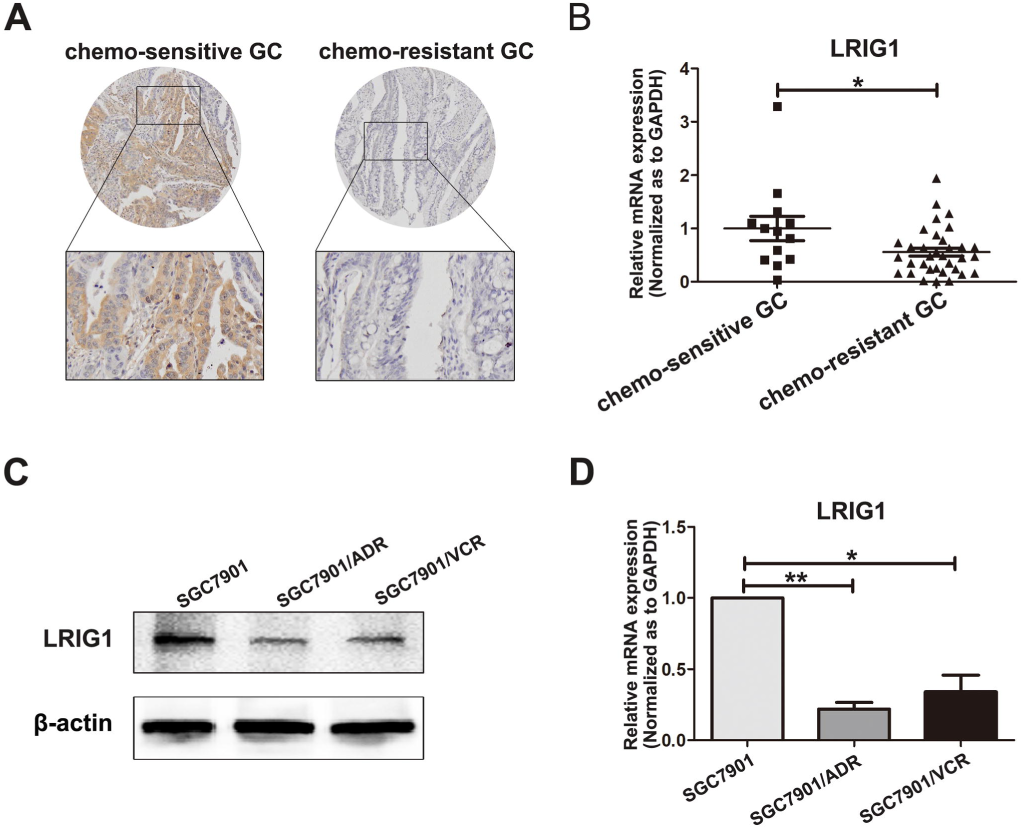
Decreased LRIG1 expression is associated with chemo-resistance in GC. (A) Immunohistochemistry analysis of LRIG1 expression in chemo-sensitive and chemo-resistant GC tissues. (B) The expression level of LRIG1 mRNA in chemo-sensitive and chemo-resistant GC tissues was measured using qRT-PCR. GAPDH was used as an internal control and the fold change was calculated by 2^-ΔΔCt^. (C) The expression of LRIG1 in GC cell line SGC7901 and its MDR variants SGC7901/VCR and SGC7901/ADR was examined through western blotting analysis. β-actin was used as an internal control. (D) The expression level of LRIG mRNA in GC cell line SGC7901 and its MDR variants SGC7901/VCR and SGC7901/ADR was measured using qRT-PCR. GAPDH was used as an internal control.

**Table 1.** LRIG1 expression in chemo-sensitive and chemo-resistant gastric cancer tissues

|  | n | Expression level of LRIG1 |  |  |  | *P* |
| --- | --- | --- | --- | --- | --- | --- |
|  |  | – | + | ++ | +++ |  |
| **chemo-sensitive GC** | 13 | 4 | 2 | 3 | 4 | <0.05 |
| **chemo-resistant GC** | 34 | 19 | 9 | 5 | 1 |  |

$\chi^2$ test were used to evaluate the significance of differences in two groups.

### Up-regulation of LRIG1 enhances chemo-sensitivity of GC cells

Next, we investigated whether up-regulation of LRIG1 increased sensitivity to chemotherapeutic agents in GC cells. To this end, SGC7901 cells were transfected with siRNA against LRIG1 (siLRIG1) or NC siRNA (siNC), and SGC7901/ADR and SGC7901/VCR cells were transfected with LRIG1 vector or NC (negative control). Western blotting assay confirmed that siLRIG1 dramatically suppressed LRIG1 expression in SGC7901, while LRIG1 vector markedly up-regulated LRIG1 expression in SGC7901/ADR and SGC7901/VCR (Figure 3A). We found that silencing of LRIG1 significantly inhibited sensitivity to chemotherapy, as indicated by an increase in the IC50 values of VCR (vincristine), ADR (adriamycin), 5-FU (5-fluorouracil) and CDDP (cisplatin) in SGC7901 cells (Figure 3B). Meanwhile, deceased IC50 values for these four chemotherapeutic agents were observed after LRIG1 expression was up-regulated by LRIG1 vector in SGC7901/ADR and SGC7901/VCR cells (Figure 3C and 3D). Since one major mechanism through which tumor cells escape from drug-induced damage is alterations in drug influx and efflux, we analyzed intracellular accumulation and release of ADR. As shown in Table 2, down-regulation of LRIG1 in SGC7901 cells manifested an increased releasing index for ADR compared with SGC7901-siNC cells. Conversely, restoration of LRIG1 in SGC7901/ADR and SGC7901/VCR cells showed a diminished releasing index for ADR. Moreover, the function of LRIG1 in GC cell MDR was also verified by the increase of apoptosis induced by 5-FU in SGC7901/ADR and SGC7901/VCR cells after up-regulation of LRIG1. In contrast, reduced LRIG1 expression in SGC7901 cells led to a decrease in the apoptosis (Figure 3E). Taken together, these results suggested that LRIG1 correlates with GC MDR and up-regulation of LRIG1 could increase sensitivity to chemotherapeutic drugs.

**Figure 3.**
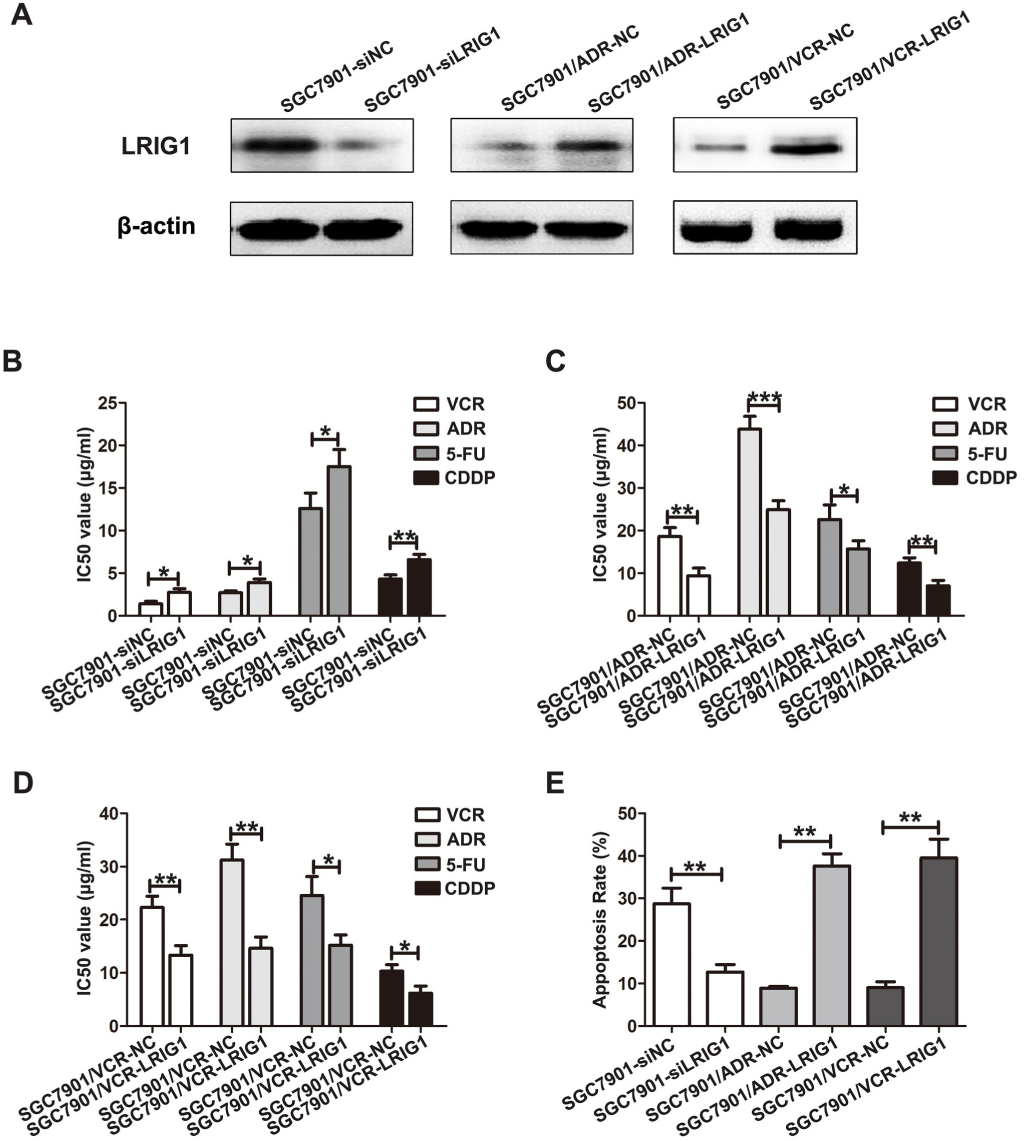
Up-regulation of LRIG1 enhances chemo-sensitivity of GC cells. (A) Western blotting analysis of LRIG1 expression in SGC7901 cells transfected with LRIG1 siRNA (siLRIG1) or negative control (siNC), and SGC7901/ADR and SGC7901/VCR cells transfected with LRIG1 plasmid or vector control (NC). β-actin was used as an internal control. (B) IC50 values of SGC7901 cells to VCR, ADR, 5-FU, and CDDP calculated from MTT assays after transfection with siLRIG1 or siNC. (C) IC50 values of SGC7901/ADR cells to VCR, ADR, 5-FU, and CDDP calculated from MTT assays after transfection with LRIG1 or NC. (D) IC50 values of SGC7901/VCR cells to VCR, ADR, 5-FU, and CDDP calculated from MTT assays after transfection with LRIG1 or NC. (E) The apoptotic rates of cells treated with 5-FU (50 mg/ml for SGC7901/ADR and SGC7901/VCR; 20 mg/ml for SGC7901) for 24 h were measured by flow cytometry.

**Table 2.** Intracellular ADR accumulation and retention in gastric cancer cells

| Cell lines | Fluorescence intensity |  | Releasing index | *P* |
| --- | --- | --- | --- | --- |
|  | Accumulation | Retention |  |  |
| SGC7901-siNC | 2.82±0.20 | 2.08±0.18 | 0.26±0.06 | <0.05 |
| SGC7901-siLRIG1 | 1.41±0.16 | 0.87±0.09 | 0.38±0.04 |  |
| SGC7901/VCR-NC | 2.01±0.18 | 1.09±0.13 | 0.46±0.06 | <0.05 |
| SGC7901/VCR-LRIG1 | 3.08±0.31 | 2.05±0.24 | 0.33±0.03 |  |
| SGC7901/ADR-NC | 1.94±0.13 | 1.09±0.13 | 0.44±0.05 | <0.01 |
| SGC7901/ADR-LRIG1 | 3.63±0.29 | 2.85±0.17 | 0.21±0.03 |  |
| SGC7901-mimic NC | 2.75±0.14 | 1.99±0.15 | 0.28±0.07 | <0.05 |
| SGC7901-miR-20a mimic | 1.29±0.08 | 0.72±0.10 | 0.44±0.06 |  |
| SGC7901/VCR-inhibitor NC | 2.04±0.20 | 1.11±0.11 | 0.46±0.08 | <0.05 |
| SGC7901/VCR-miR-20a inhibitor | 3.16±0.29 | 2.25±0.18 | 0.29±0.03 |  |
| SGC7901/ADR- inhibitor NC | 2.01±0.23 | 1.14±0.14 | 0.43±0.10 | <0.05 |
| SGC7901/ADR- miR-20a inhibitor | 3.41±0.31 | 2.82±0.20 | 0.17±0.06 |  |

### miR-20a down-regulates LRIG1 expression by binding its 3′-UTR

To explore the regulatory mechanism of LRIG1 at miRNA level, we employed bioinformatic methods to identify the potential miRNAs regulating LRIG1. Based on the bioinformatics analysis of three different programs (miRanda, Pictar and targetscan), a highly conserved miR-20a targeting sequence was predicted in the 3′-UTR of the LRIG1 mRNA. We then examined the relationship between the expression levels of LRIG1 mRNA and miR-20a in the GC drug resistant cell lines. Compared with the parent cell line SGC7901, the expression of miR-20a was markedly higher in the resistant cell lines SGC7901/ADR and SGC7901/VCR (Figure 4A), inverse with the expression of LRIG1 in the GC cell lines. To further address whether miR-20a directly targeted LRIG1 3′-UTR to inhibit its expression in GC, PCR products containing wildtype or mutant LRIG1 3′-UTR binding sites were inserted into reporter vectors (Figure 4B). Luciferase reporter assays showed that miR-20a brought a significant reduction in relative luciferase activity when the LRIG1 plasmid containing wild-type 3′-UTR was present, but the luciferase activity was not significantly changed in the 3′-UTR with mutant binding sites (Figure 4C). This result indicated that miR-20a might repress LRIG1 expression by targeting the binding sites of the LRIG1 3′-UTR. Moreover, both qRT-PCR and Western blotting analysis showed that up-regulation of miR-20a markedly decreased LRIG1 expression in SGC7901 cells and silencing of miR-20a dramatically raised LRIG1 expression in SGC7901/ADR and SGC7901/VCR cells (Figure 4D and 4E). Collectively, these results indicated that miR-20a suppresses LRIG1 expression by directly targeting its 3′-UTR.

**Figure 4.**
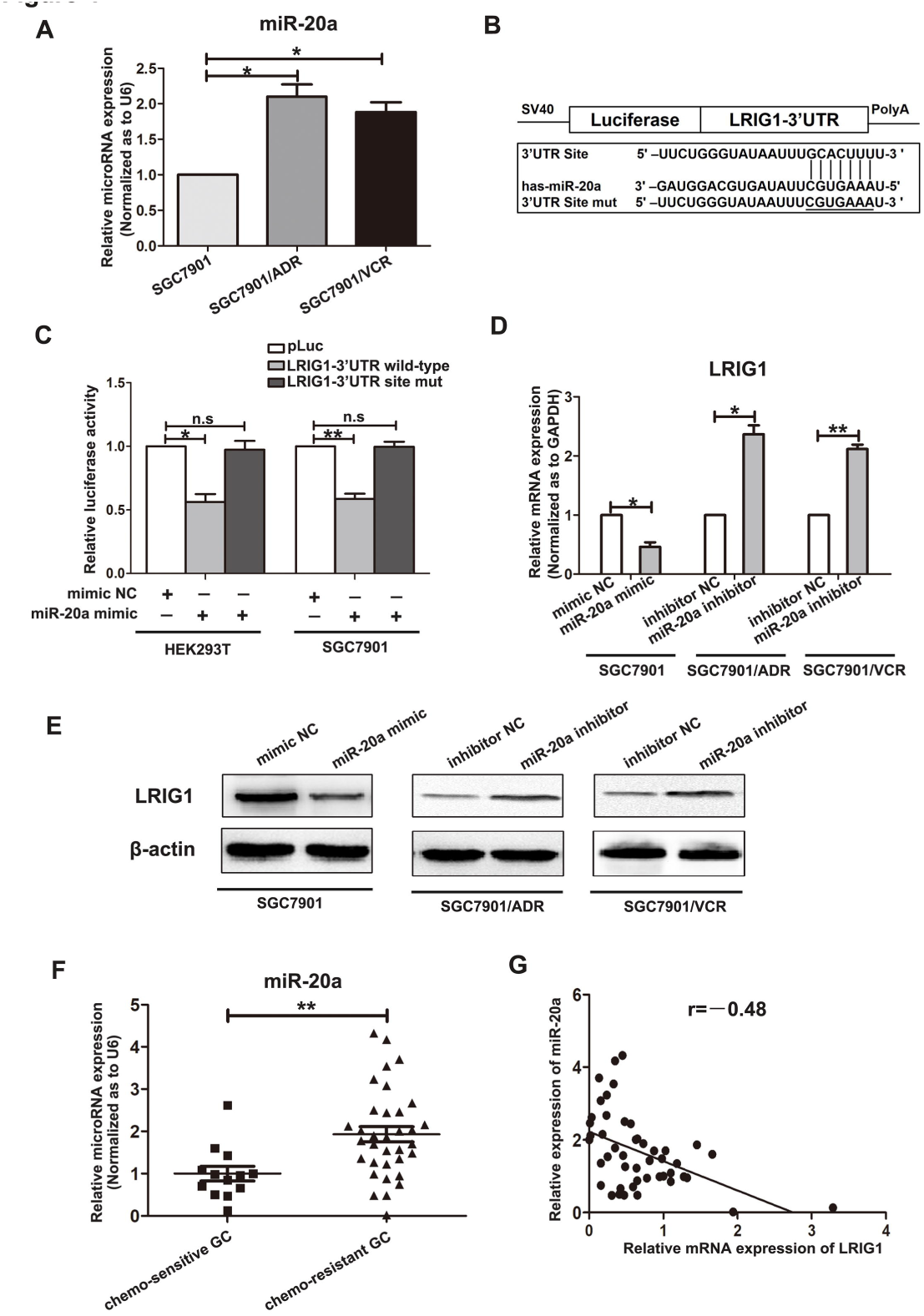
miR-20a down-regulates LRIG1 expression by binding its 3′-UTR. (A) The expression level of miR-20a in GC cell line SGC7901 and its MDR variants SGC7901/VCR and SGC7901/ADR. The U6 small nuclear RNA was used as an internal control and the fold-change was calculated using 2^-ΔΔCt^. (B) Diagram of LRIG1 3′-UTR-containing reporter construct. Mutations were generated at the predicted miR-20a binding site located in the LRIG1 3′-UTR. (C) The luciferase activity after the wild type or mutant reporter plasmids were co-transfected with miR-20a mimic or NC in HEK293T and SGC7901 cells. (D) The expression of LRIG1 mRNA in SGC7901 cell after transfection with miR-20a mimic or mimic control (mimic NC), and SGC7901/ADR and SGC7901/VCR cells transfected with miR-20a inhibitors or inhibitor control (inhibitor NC) was analysed using qRT–PCR. GAPDH was used as an internal control. (E) The expression of LRIG1 protein was analysed in SGC7901 cell after transfection with miR-20a mimic or mimic NC, and SGC7901/ADR and SGC7901/VCR cells transfected with miR-20a inhibitors or inhibitor NC through western blotting. β-actin served as an internal control. (F) The expression of miR-20a in chemo-sensitive and chemo-resistant GC tissues was examined by qRT-PCR. U6 was used as an internal control. (G) A statistically significant inverse correlation between miR-20a and LRIG1 mRNA levels in GC specimens (Spearman’s correlation analysis, r=-0.48; P < 0.01).

We next measured whether miR-20a expression was associated with LRIG1 expression in GC samples to evaluate the clinical relevance of miR-20a/LRIG1 axis. The expression of miR-20a in GC tissues from 47 GC patients was detected by qRT-PCR. We found miR-20a was significantly overexpressed in chemo-resistant GC tissues, compared with chemo-sensitive cases (Figure 4F). Statistical analysis revealed that LRIG1 expression was inversely correlated with miR-20a expression (Figure 4G). These observations suggested that miR-20a and LRIG1 are inversely expressed in GC specimens.

### miR-20a is involved in chemo-resistance of GC cells

To verify the effect of miR-20a on GC drug resistance, we transfected miR-20a mimic into SGC7901 cells, and miR-20a inhibitor into SGC7901/ADR and SGC7901/VCR cells, to conduct gain- and loss-of-function experiments. Remarkably, the restoration of miR-20a in SGC7901 cells increased the IC50 values to VCR, ADR, 5FU and CDDP (Figure 5A), and exhibited an increased releasing index for ADR (Table 2). Meanwhile, up-regulation of miR-20a led to reduced apoptosis (Figure 5D). In contrast, inhibition of miR-20a in SGC7901/ADR and SGC7901/VCR cells sensitized drug-resistant cells to chemotherapeutics by increasing the intracellular concentration of ADR (Table 2). Silencing of miR-20a decreased the IC50 values and also markedly raised apoptosis rates (Figure 5B, 5C and 5D). Therefore, these results indicated that miR-20a has a negative effect on GC chemo-sensitivity.

**Figure 5.**
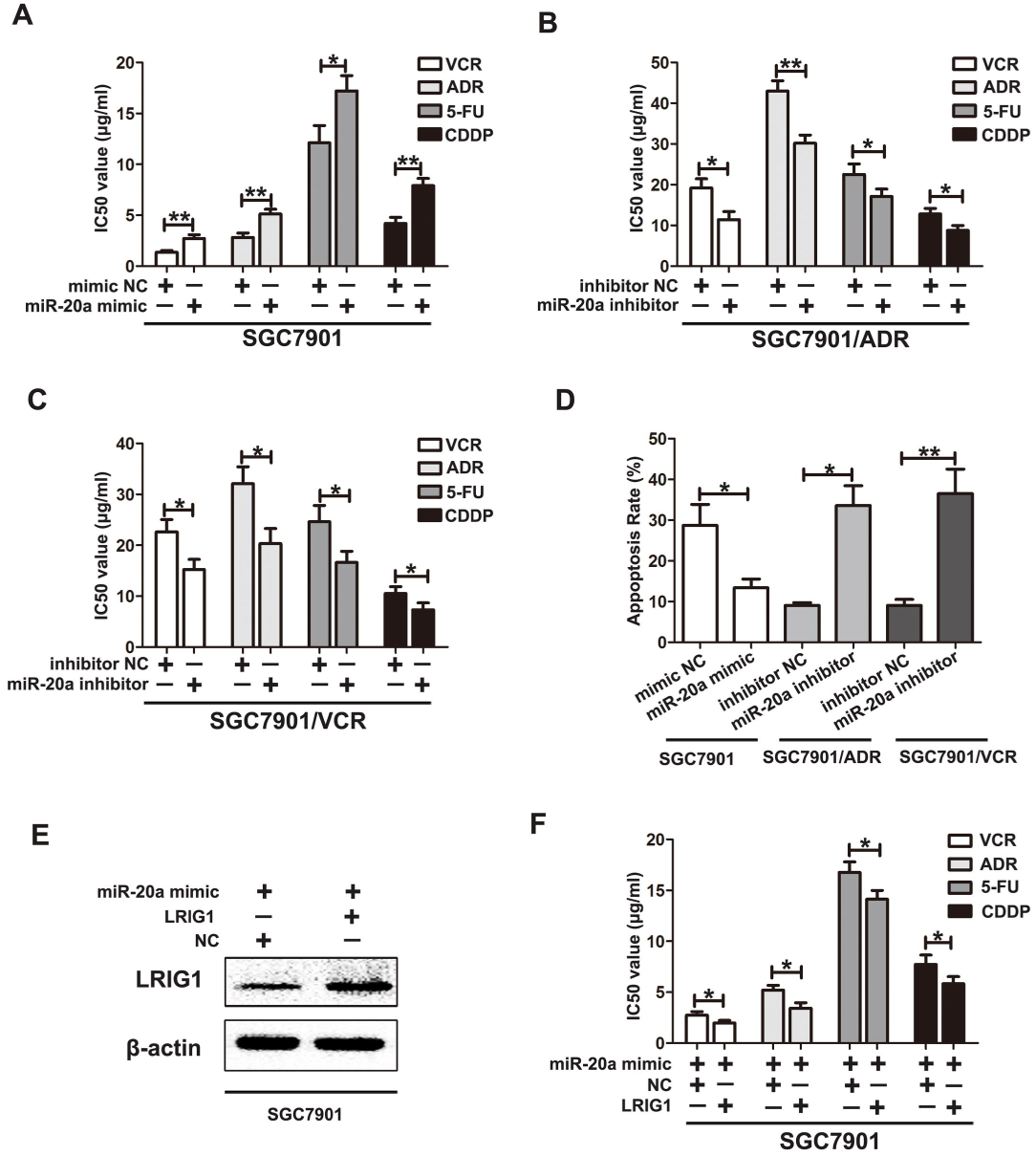
miR-20a is involved in chemo-resistance of GC cells. (A) IC50 values of SGC7901 cells to VCR, ADR, 5-FU, and CDDP calculated from MTT assays after transfection with miR-20a mimic or control. (B) IC50 values of SGC7901/ADR cells to VCR, ADR, 5-FU, and CDDP calculated from MTT assays after transfection with miR-20a inhibitor or control. (C) IC50 values of SGC7901/VCR cells to VCR, ADR, 5-FU, and CDDP calculated from MTT assays after transfection with miR-20a inhibitor or control. (D) The apoptotic rates of cells treated with 5-FU for 24 h were measured by flow cytometry. (E) Western blotting of LRIG1 in SGC7901 cells co-transfected with miR-20a mimic and LRIG1 plasmid. β-actin was used as an internal control. (F) IC50 values of SGC7901 cells to VCR, ADR, 5-FU, and CDDP calculated from MTT assays after co-transfection with miR-20a mimic and LRIG1 plasmid.

Since LRIG1 can inhibit GC cell chemo-resistance and miR-20a can post-transcriptionally regulate the expression of LRIG1, we hypothesized that miR-20a mediated down-regulation of LRIG1 directly promotes GC chemo-resistance. To address this hypothesis, we first transfected SGC7901 cells with LRIG1 plasmids, and then miR-20a mimic was co-transfected into SGC7901 cells (Figure 5E). *In vitro* drug sensitivity assay demonstrated that the restoration of miR-20a expression significantly reduced LRIG1-induced GC cell chemo-sensitivity (Figure 5F).

### miR-20a/LRIG1 axis regulates GC drug resistance through EGFR mediated PI3K/AKT and MAPK/ERK signaling

It has been demonstrated that LRIG1 is capable of interacting with EGFR and enhancing both its basal and ligand-stimulated ubiquitination and degradation(Gur et al., 2004; Powell et al., 2012; Wang et al., 2013). Thus, we speculated that LRIG1 modulated GC drug resistance through EGFR signaling. qRT-PCR and western blotting assays were performed to investigate whether EGFR activity was changed after LRIG1 expression altered. It was shown that down-regulation of LRIG1 in SGC7901 cells did not result in an impact on the mRNA level of EGFR, but led to a substantial increase in the protein level of EGFR (Figure 6A and 6B). It can be inferred that LRIG1 might directly interact with EGFR protein, but not via transcription regulation. Subsequently, a co-immunoprecipitation method was carried out to determine whether there was a physical interaction between LRIG1 and EGFR molecules. As shown in Figure 6C, EGFR could be specifically co-immunoprecipitated with LRIG1, but not with control IgG, indicating that these two proteins were specifically associated with each other in complex. In addition, EGFR expression was detected through IHC in the 47 GC tissues, and statistical analysis revealed that LRIG1 expression was significantly negatively correlated with EGFR levels (Table 3).

**Figure 6.**
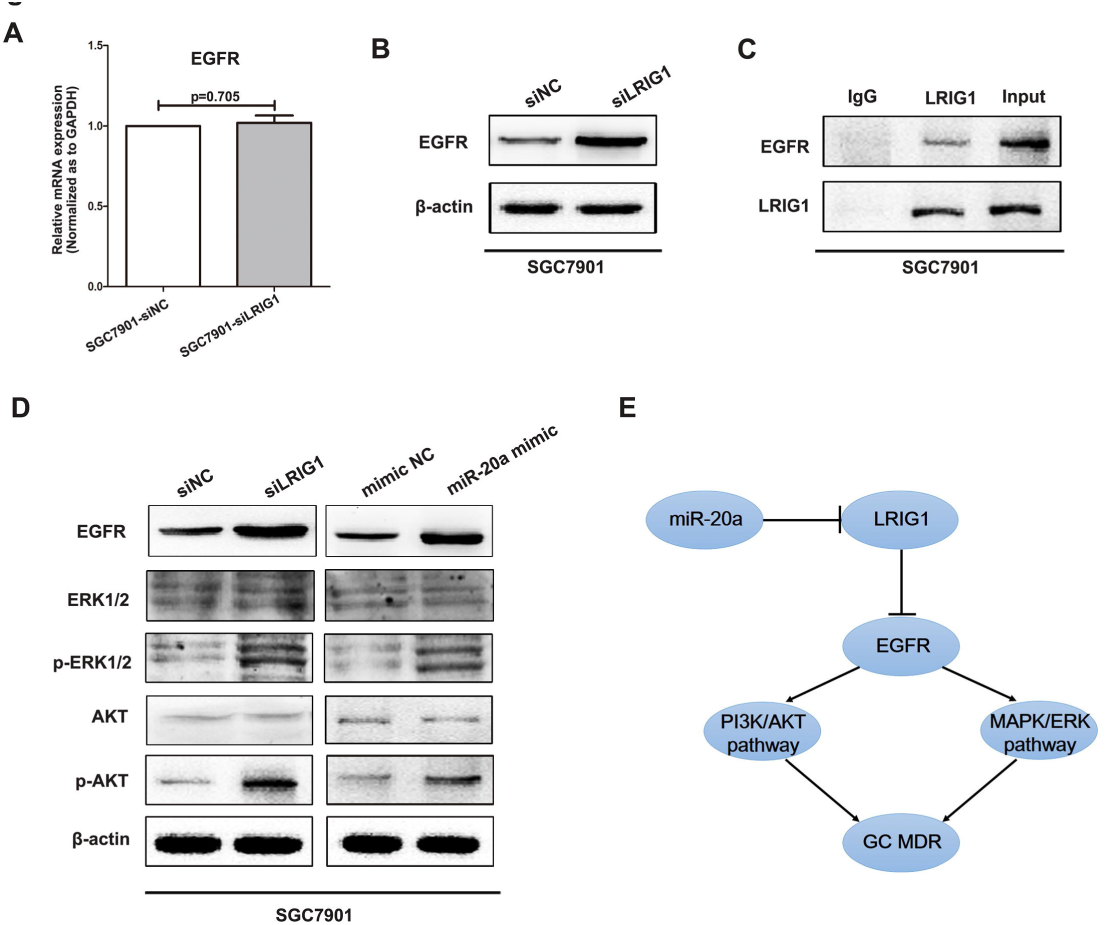
miR-20a/LRIG1 axis regulates GC chemo-resistance through EGFR mediated PI3K/AKT and MAPK/ERK signaling. (A) The mRNA expression of EGFR in SGC7901 was examined by qRT-PCR after transfection with siLRIG1 or siNC. GAPDH was used as an internal control. (B) Western blotting of LRIG1 in SGC7901 cells transfected with siLRIG1 or siNC. β-actin was used as an internal control. (C) Lysates of SGC7901 cells were immunoprecipitated with anti-LRIG1 or control IgG and blotted with antibodies to EGFR or LRIG1. (D) Western blotting of ERK1/2, AKT and their phosphorylated forms in SGC7901 cells after transfection with siLRIG1 or miR-20a mimic. β-actin was used as an internal control. (E) A schematic representation of miR-20a/LRIG1 axis as a MDR regulator in GC.

**Table 3.** The correlation between LRIG1 and EGFR in gastric cancer tissues

|  |  | Expression level of LRIG1 |  |  |  | Spearman's test |  |
| --- | --- | --- | --- | --- | --- | --- | --- |
|  |  | – | + | ++ | +++ | **r** | **P** |
| **Expression**<br>**level**<br>**of**<br>**EGFR** | – | 4 | 2 | 3 | 3 | –0.299 | <0.05 |
|  | + | 5 | 3 | 2 | 1 |  |  |
|  | ++ | 7 | 3 | 2 | 1 |  |  |
|  | +++ | 7 | 3 | 1 | 0 |  |  |

It is well known that EGFR mediated PI3K/AKT and MAPK/ERK pathways play very important roles in GC drug resistance(Liang et al., 2009; Roskoski, 2014; Sun et al., 2014; Yu et al., 2008). Therefore, we next measured the expression alteration of key molecules in PI3K/AKT and MAPK/ERK pathways to further clarify the miR-20a/LRIG1 downstream mechanism. Western blotting demonstrated that down-regulation of LRIG1 or up-regulation of miR-20a significantly enhanced EGFR activity, and increased p-AKT (phospho-AKT) and p-ERK1/2 (phospho-ERK1/2) protein expression (Figure 6D). Taken together, these results indicated that miR-20a/LRIG1 axis might regulate GC cell drug resistance by increasing EGFR activity and secondary accelerating PI3K/AKT and MAPK/ERK signaling pathways (Figure 6E).

## Discussion

MDR accounts for most cases of treatment failure among GC patients(Van Cutsem et al., 2016). Although a wide range of MDR-associated molecules have been discovered, the mechanisms underlying their functions and their interactions are not fully understood. The human LRIG1 gene at chromosome 3p14.1 encodes LRIG1 protein that negatively regulates the ERRB2 gene product, human epidermal growth factor receptor 2 (HER2), and other oncogenic receptor tyrosine kinases, including EGFR, HER3 and HER4 (Gur et al., 2004; Miller et al., 2008; Morrison et al., 2016; Powell et al., 2012; Shattuck et al., 2007; Wang et al., 2013). Recently, LRIG1 has been shown to function as a tumor suppressor in the mouse intestine(Powell et al., 2012), and increasing evidence indicates that LRIG1 may act as a tumor suppressor in humans(Zhu et al., 2010). Reduced expression of LRIG1 has been reported in breast, cervical, and skin cancers, as reviewed (Hedman and Henriksson, 2007; Wang et al., 2013). The soluble ectodomain of LRIG1 inhibits *in vivo* growth of EGFRVIII mutant gliomas, and restoration of LRIG1 expression sensitizes glioma cells to chemotherapy(Stutz et al., 2008). In the present study, we observed that LRIG1 was down-regulated in GC tissues, associated with chemo-sensitivity, and low expression of LRIG1 predicted poor prognosis in GC individuals. Moreover, LRIG1 correlated with MDR of GC and up-regulation of LRIG1 could increase sensitivity to chemotherapeutic drugs.

Accumulating evidence indicates that miRNAs play important roles in the MDR of various cancers. miRNAs, including miR-27a, miR-497, miR-363, miR-181b, miR-15b and miR-16, have been reported to be implicated in the MDR of GC(Ishimoto et al., 2016; Riquelme et al., 2016; Song and Ajani, 2013; Xia et al., 2008; Zhang et al., 2016; Zhu et al., 2010). Our previous studies identified 11 miRNAs, which regulate MDR in GC, using high-throughput functional screening, and further confirmed that miR-27b/CCNG1/P53/miR-508-5p axis plays important roles in GC-associated MDR(Shang et al., 2016; Shang et al., 2014). Here we identified miR-20a as an upstream miRNA of LRIG1, playing an important role in GC drug resistance. miR-20a suppressed LRIG1 expression by directly targeting its 3′-UTR, and was inversely expressed with LRIG1 in GC specimens. In addition, restoration of miR-20a expression significantly reduced LRIG1-induced GC cell chemo-sensitivity.

miR-20a, a member of miR-17-92 cluster, has already been widely studied in different cancers, acting as an oncogene in tumor development and progression(Chang et al., 2013; Liu et al., 2017; Wang et al., 2015; Weng et al., 2014). It was reported that ectopic expression of miR-20a could lead to epithelial-mesenchymal transition and promote metastasis of gallbladder carcinoma, by directly binding the 3′-UTR of Smad7, and subsequently enhancing nuclear translocation of β-catenin(Chang et al., 2013). More recently, Liu *et al.*(Liu et al., 2017) demonstrated that miR-20a-mediated autophagy defect might be a new mechanism underlying the oncogenic function of miRNA during breast tumorigenesis. Furthermore, miR-20a was shown to modulate sensitivity to some anti-cancer drugs, such as the anti-leukemia drug etoposide (VP-16), 5-FU, oxaliplatin (L-OHP), and teniposide (VM-26). For example, suppression of miR-20a expression by treatment with either miR-20a inhibitor or oridonin sensitized leukemia cells to VP-16(Weng et al., 2014). It was also reported that miR-20a regulated chemotherapeutic sensitivity of colorectal adenocarcinoma cancer cells to the most used cancer chemotherapy agents(Chai et al., 2011). Furthermore, miR-20a enhanced cisplatin resistance of GC via targeting cylindromatosis(Zhu et al., 2016). This study further confirmed that miR-20a is involved in MDR of GC, and acts as an oncogenic miRNA.

Currently, the tumor suppressive function of LRIG1 is mainly attributed to its inhibitory effect on EGFR signaling(Gur et al., 2004; Powell et al., 2012; Wang et al., 2013). Over the past few years, it has been shown that LRIG1 binds to EGFR and attenuates EGFR signaling via both receptor degradation and catalytic inhibition(Gur et al., 2004; Powell et al., 2012; Sheu et al., 2014; Wang et al., 2013). EGFR is a well-studied, versatile signal transducer which is overexpressed in many types of tumor cells, including lung, colon and prostatic carcinoma, and up-regulation of EGFR is associated with poor clinical prognosis(Burgess et al., 2003; Guo et al., 2015; Roskoski, 2014). EGFR mediates signals that stimulate proliferation, migration, and metastasis in many tumor types, and its signal transduction is regulated by stimulatory and inhibitory inputs(Lo and Hung, 2006; Roskoski, 2014). Yu *et al*.(Yu et al., 2008) demonstrated that PI3K/AKT pathway could be inactivated by doxorubicin and etoposide, and wortmannin could increase the sensitivity of GC cells to chemotherapy. Zhang *et al*.(Zhang et al., 2017) demonstrated that miR-939 contributed to chemo-sensitivity by inhibiting SLC34A2/ Raf/MEK/ERK pathway. The present study showed that miR-20a/LRIG1 axis might regulate GC cell drug resistance by increasing EGFR activity and secondary promotion of PI3K/AKT and MAPK/ERK signaling pathways.

In conclusion, the results obtained in the present study reveal that low expression of LRIG1 in GC drug resistance might reflect the increased levels of miR-20a, which could significantly promote MDR in GC. Furthermore, miR-20a/LRIG1 axis might regulate GC drug resistance through EGFR mediated PI3K/AKT and MAPK/ERK signaling. The newly identified miR-20a/LRIG1/EGFR link provides a new potential therapeutic strategy for the future treatment of GC patients with MDR.

## Material and methods

### Cell culture and tissue collection

The human GC cell line SGC7901 and its multidrug-resistant variants SGC7901/VCR and SGC7901/ADR were obtained from State Key Laboratory of Cancer Biology and Xijing Hospital of Digestive Diseases, and maintained in RPMI-1640 medium (Hyclone, Logan, UT, USA) supplemented with 10% fetal bovine serum (Gibco, Carlsbad, CA, USA) at 37°C under a humidified air atmosphere containing 5% CO_2_. Paired samples of primary GC and adjacent normal tissues were obtained from patients who had undergone surgery for GC resection at Xijing Hospital, Xi’an, China. This study was approved by the Protection of Human Subjects Committee of our hospital, and informed consent was obtained from each patient.

### Immunohistochemistry

IHC was performed as previous study(Zhou et al., 2014) with an anti-LRIG1 (Abcam, Cambridge, UK) and anti-EGFR (Cell Signaling, Danvers, MA, USA) antibodies. The protein expression was visualized and classified according to the percentage of positive cells and the intensity of staining, and the histological score (H score) was calculated and graded as described in our previous study(Zhou et al., 2014).

### RNA extraction and real-time quantitative reverse transcription–polymerase chain reaction (qRT-PCR)

Total RNA was extracted with Trizol reagent (Invitrogen, Carlsbad, CA, USA). The RT and PCR primers for miR-20a and U6 were purchased from RiBoBio (Guangzhou, China). The PCR primers for LRIG1 and EGFR are shown in Supplementary Table 1. The first-strand cDNA was synthesized with the PrimeScript RT reagent Kit (Takara, Dalian, China). RT-PCR was performed with SYBR Premix Ex TaqII (Takara) and measured in the LightCycler 480 (Roche, Basel, Switzerland). Glyceraldehyde 3-phosphate dehydrogenase (GAPDH) or U6 snRNA was used as an endogenous control, and an internal control was used to verify that sample loading was equal. The fold change was calculated by 2^-ΔΔCt^. All reactions were performed in triplicate.

### Western blotting

Whole-cell lysates were prepared in RIPA buffer (Byotime, Haimen, China), and Western blotting analysis was performed(Zhou et al., 2014). The primary antibodies used were anti-LRIG1 (Abcam), anti-EGFR (Cell Signaling), anti-ERK1/2 (Abcam), anti-p-ERK1/2 (Abcam), anti-AKT (Cell Signaling), anti-p-AKT (Cell Signaling) and β-actin (Sigma, St. Louis, MO, USA).

#### Oligonucleotide construction and transfection

Mimics and inhibitors for has-miR-20a, and negative control oligonucleotides were purchased from RiboBio. siRNAs and plasmids for LRIG1, and corresponding negative control were obtained from Genechem Co., Ltd. (Shanghai, China). Target cells were transfected with oligonucleotides using Lipofectamine 2000 reagent (Invitrogen) according to the manufacturer’s instructions.

#### *In vitro* drug sensitivity assay

Drug sensitivity was assessed as described previously(Shang et al., 2014). Briefly, 5×10^3^ cells were seeded in 96-well plates, and medium containing chemotherapeutic drugs was added to each well. After incubation for 48 h, a 3-(4,5-di-methyl-2-thiazolyl)-2,5- diphenyl-2H tetrazolium bromide (MTT, Sigma) assay was performed. Inhibition rates and IC50 values were then calculated.

#### Apoptosis assay

Cell apoptosis was evaluated using an Annexin-V-FITC apoptosis detection kit (BD, Franklin Lakes, NJ, USA) as previously described(Shang et al., 2014).

#### Analysis of intracellular ADR concentrations

Fluorescence intensity of intracellular ADR was determined by flow cytometry as described previously(Liang et al., 2009). Briefly, cells were seeded into six-well plates (1×10^6^ cells/well) and cultured for 1h after ADR addition. Cells were then either harvested to detect ADR accumulation or cultures were continued in a drug-free medium for another 2 h, followed by detection of ADR retention. The releasing index of ADR in the GC cells was calculated using the following formula: releasing index = (accumulation value – retention value)/accumulation value.

#### Luciferase assay

Plasmids carrying wild-type Luc-LRIG1 or mutant Luc-LRIG1-β3’-UTR were synthesized (GeneCopoeia, Rockville, MD, USA). The luciferase assay was performed as previously described(Zhou et al., 2013).

#### Immunoprecipitation

Immunoprecipitation assay was performed as previously described using an anti-LRIG1 antibody(Dong et al., 2017). The total protein was prepared using M-PERTM mammalian protein extraction reagent (Pierce, Appleton, WI, USA). 10% of chromatin was used as an input control, and a non-specific antibody (anti-IgG, Abcam) served as a negative control. The obtained proteins were subjected to Western blotting in an attempt to amplify the LRIG1-binding sites.

### Statistical analysis

SPSS software (version 21.0, SPSS Inc., Chicago, IL, USA) was used for the statistical analyses. The continuous data were presented as the means ± standard errors of the mean, and compared using Student’s t-test (two-tailed) or oneway analysis of variance (ANOVA). The Spearman’s correlation test was performed to examine the relationship of LRIG1 and miR-20a or EGFR expression in GC tissues. P<0.05 was considered statistically significant (*P<0.05, **P<0.01 and ***P<0.001).

## Acknowledgements

We would like to thank Zheng Chen and Jianhua Dou from the State Key Laboratory of Cancer Biology for the assistance in tissue sample collection.

## Abbreviations

5-FU: 5-fluorouracil
ABCB1: ATP-binding cassette, sub-family B member 1
ABCG2: ATP-binding cassette, sub-family G, member2
ADR: adriamycin
AKT: protein kinase B
CDDP: cisplatin
EGFR: epidermal growth factor receptor
ERK: extracellular signal-regulated kinase
GAPDH: glyceraldehyde 3-phosphate dehydrogenase
GC: gastric cancer
HER2: human epidermal growth factor receptor 2
IHC: immunohistochemistry
LRIG1: leucine-rich repeats and immunoglobulin-like domains 1
MAPK: mitogen-activated protein kinase
MDR: multidrug resistance
miR: microRNA
miRNA: microRNA
mRNA: messenger RNA
PI3K: phosphatidylinositol 3 kinase
qRT-PCR: quantitative reverse transcription–polymerase chain reaction
siRNA: silencing RNA
TNFα: tumor necrosis factor α
UTR: untranslated region
VCR: vincristine

